# Enzyme Classification via Semi-Supervised Functional Residue Learning

**DOI:** 10.64898/2026.02.11.705200

**Authors:** Chengyue Gong, Daiheng Zhang, Jeffrey Ouyang-Zhang, Qiang Liu, Adam Klivans, Daniel Diaz

## Abstract

Predicting enzymatic function from a protein sequence is a fundamental task in protein discovery and engineering. In this paper, we present **S**emi-supervised **L**earning for **E**nzym**e C**lassification (SLEEC): a semi-supervised learning framework that learns a function-aware protein representation for Enzyme Commision (EC) number prediction. SLEEC achieves SOTA performance on standard benchmarks and provides interpretable, residue-level annotations. We further demonstrate that our framework is robust to benign sequence modifications routinely observed in protein engineering workflows– such as appending functional tags– a desirable property that current ML frameworks lack. Our main technical contribution is a multiple sequence alignment (MSA)-based data augmentation technique for discovering sparse residue activations within a given enzyme sequence.

## 1 Introduction

Understanding the sequence-function relationship of proteins is critical to our discovery and development of biotechnology for industrial and pharmaceutical applications. Experimental characterization of protein function is too labor-intensive to keep up with the exploding number of discovered protein sequences. As of 2024, we sequence over 100K new proteins a day and have cumulatively discovered over 248 million unique sequences [8], but only 0.2% (∼ 570,000) of discovered sequences have experimental characterization [4, 8]. Homology-based computational approaches have enabled the putative assignment of enzymatic function but often misannotate enzymes that have mutations in functionally relevant motifs or fail to annotate sequences that lack an evolutionary relationship with experimentally characterized proteins [*e*.*g*., 1, 2, 37, 43, 45, 53, 56].

Recently, machine learning methods have shown promise in computational function annotation of protein sequences, specifically the prediction of Enzyme Commission (EC) number, a classification ontology for enzymatic function [*e*.*g*., 9, 23, 38, 54]. These methods commonly overfit to the sequence dataset and demonstrate poor generalization on held-out validation sets. More importantly, an often overlooked drawback of current ML frameworks is their lack of robustness to sequence modifications that do not affect function but are routinely observed in protein discovery and engineering workflows. For example, functional tags are often appended to the N- or C-terminus of a protein to improve expression, trigger secretion or localization to a cellular compartment, and do not modify enzymatic function. During protein engineering, orthologs– functionally related proteins that differ due to speciation– are often screened in parallel to select a starting point to engineer. In these scenarios, the existing state-of-the-art (SOTA) framework, CLEAN [54], demonstrates poor robustness (see Table 1 for details). Furthermore, saliency analysis using a discriminative localization technique [57] reveals that CLEAN’s predictions are evenly weighted across the sequence, rather than being driven by the subset of residues responsible for substrate specificity and catalysis (see Figure 1). This example underscores the need for developing ML frameworks for EC number prediction that are robust to routine sequence modifications and offer residue-level interpretability.

**Table 1:** CLEAN is not robust to common sequence modifications encountered in protein engineering workflows. We observe worse performance on sequences modified with functional tags or replaced with an ortholog. Here, we evaluate appending the Halo-(297mer) and TAP-(72mer) tags to the N-terminus of proteins in the test sets. Ortholog indicates a homologous sequence with ∼80% sequence similarity to the proteins in the test sets. ‘Sensitivity’ is the % difference in performance compared to the original sequence in the test sets.

| Metric | New-392 |  |  |  | Price-149 |  |  |  |
| --- | --- | --- | --- | --- | --- | --- | --- | --- |
|  | Original | Halo-Tag | TAP-Tag | Ortholog | Original | Halo-Tag | TAP-Tag | Ortholog |
| F1 ↑ | 0.497 | 0.302 | 0.434 | 0.405 | 0.495 | 0.190 | 0.342 | 0.412 |
| Sensitivity ↓ | 0% | 39% | 13% | 19% | 0% | 62% | 31% | 17% |

**Figure 1.**
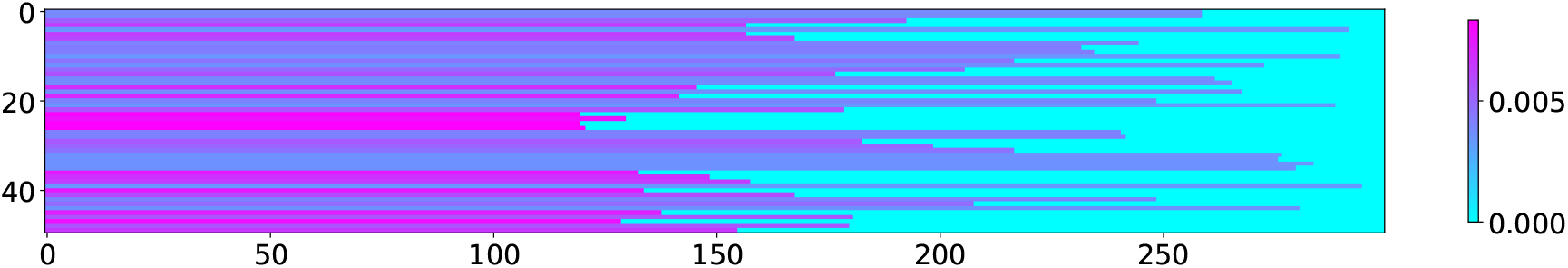
Saliency analysis of CLEAN. Using a discriminative localization technique [57] on CLEAN, we visualize saliency map values for 50 randomly selected proteins from New-392. The x-axis corresponds to sequence position and the y-axis corresponds to the 50 proteins. The cyan color indicates padding the sequence to a length of 300. The saliency analysis demonstrates that CLEAN’s predictions have a uniform weighting across the sequence and are not driven by functional residues.

In this paper, we present **S**emi-supervised **L**earning for **E**nzym**e C**lassification (SLEEC): a two-stage sequence-based framework for EC number classification. SLEEC learns a *function-aware protein representation* in order to predict the EC numbers and provide residue-level interpretability. To learn function-aware protein representations, a large training set of labeled functional residues is essential. However, datasets of experimentally determined functional residues are scarce, with the Mechanism and Catalytic Site Atlas (mCSA) [34] containing only about 4,000 functionally annotated residues across roughly 1,000 enzymes. To address this data scarcity, we propose a semi-supervised pre-training approach that combines mCSA data with evolutionary priors encoded in multiple sequence alignments of proteins [*e*.*g*., 7, 10, 16, 26]. We evaluate the performance of SLEEC on well-established benchmarks for predicting EC numbers: New-392 and Price-149. On New-392, our model achieves a ∼30% improvement in F1 score compared to the next best model, CLEAN (F1 score 0.644 vs 0.497). Similarly, on Price-149, a test set where ∼15% of EC numbers are absent from the training set, SLEEC still outperforms the CLEAN by ∼6% (F1 score 0.525 vs 0.494). Our results remain consistent across various sequence similarity training/test split thresholds. Finally, we demonstrate how our approach leads to predictions that are robust to routine sequence modifications encountered in protein engineering workflows.

## 2 Related Work

### 2.1 Multiple Sequence Alignments in Machine Learning

The rich evolutionary information captured in multiple sequence alignments (MSA) plays a key role in several protein-based machine learning models [*e*.*g*., 10, 14, 16, 19, 26, 29, 33, 35]. Often a model takes a protein’s MSA as input to learn a representation that capture the evolutionary covariation between residues [14, 19, 33]. In addition to serving as features, multiple sequence alignments also provide an evolutionary training signal to these models. For example, AlphaFold [19], EVE [14] and MSA-Transformer [33] predict the MSA content during training akin masked language modeling [12].

Other methods explicitly leverage the evolutionary priors present in MSAs [*e*.*g*., 10, 18, 21, 26, 30]. Intuitively, sequence positions in the MSA that are strongly conserved likely contribute to some functional or structural attribute of the protein. On the other hand, sequence positions with significant evolutionary variation not only tolerate diverse chemistries but enable adaptation to organism-specific evolutionary pressures. Profile prediction [47] is a protein language model predicting the soft label generated from the MSAs at each position. VespaG [26] is a protein language model supervised by the evolutionary scores computed analytically by GEMME [21]. COLLAPSE [10] trains a structure-based graph neural network that weights their contrastive loss by the entropy in the MSA profile. EvoRank [16], which is based on the Stability Oracle architecture [13, 17], uses the MSA-based profiles to generate rank labels for supervise training. SLEEC uses evolutionary priors from MSAs to conduct semi-supervised learning for identification of functional residues. To achieve this, we create a dataset of “pseudo-label” functional residues based on the sequence conservation observed within the MSA, thus expanding the experimentally validated functional residue dataset from the Mechanistic and Catalytic Site Atlas (mCSA), which we describe in details in Section 3.

### 2.2 Enzyme Function Classification

The goal of enzyme function classification is to predict a protein’s enzymatic function from its sequence and/or 3D structure. Most databases [3, 28, 31, 52] annotate the function of enzymes using Enzyme Commission (EC) numbers, a four-level hierarchical ontology that describes catalytic activity and substrate specificity. Currently, sequence- and structure-based homology methods are used to computationally annotate newly discovered enzymes by comparing them with their experimentally characterized evolutionary closest relatives [*e*.*g*., 1, 2, 37, 43, 45, 53, 56]. Recently, deep learning-based approaches have begun to outperform existing homology search-based approaches [*e*.*g*., 5, 9, 23, 38, 39, 46, 51, 54]. CLEAN [54] uses a contrastive learning framework that predicts the EC number of a query sequence by using cosine similarity to EC number cluster embeddings. EnzymeNet [51] and DEEPre [23] predict EC numbers through a multi-step process, where the first neural network determines the initial hierarchy level, and one of six second-level networks infers the remaining levels. EnzBert [5] appends a CLS token to the input sequence and uses a attention to implicitly learn the residues that are important for EC number prediction. SLEEC builds on this line of research but instead explicitly determines key residues to learn a function-aware representation for downstream enzyme classification. To achieve this, we use semi-supervised learning to train a residue-level binary classifier, enabling us to perform weighted pooling and obtain a function-aware representation. Quantitative comparisons can be found in Section 4.

### 2.3 Functional Residue Classification

In our work, we propose to improve EC number classification by enriching a protein’s hidden representation for functional residues. The goal of functional residue classification is to pinpoint the residues in an enzyme that are essential for its activity. These residues may be catalytic, playing a role in the reaction mechanism, or involved in determining substrate specificity. Prior work [7] has shown that functional sites are often highly conserved with destabilizing amino acids. This is known as the stability-function trade-off [27]. Traditional methods, such as PROSITE [40] and other rule-based engines [6, 25, 41], scan the protein for particular sequence or structural motifs or rules corresponding to a specific function. When the function is narrowed to substrate binding, FPocket [22] and P2Rank [20] use physics-based modeling of protein structures to identify ligand binding pockets. Recently, deep learning-based approaches have been used for functional residue classification. RXNAAMapper [49] predicts enzymatic binding site residues by first training a masked language model on SMART reactions-enzyme sequence pairs, and then identifying residues with high attention scores as binding site residues. PARSE [11] predicts whether a residue is functional by performing a nearest neighbor lookup to a database of functionally annotated residues, such as mCSA [34], in the COLLAPSE [10] latent space. In this paper, we learn a binary classifier to distinguish whether a residue is functional via semi-supervised learning.

## 3 Methods

### Overview of SLEEC

We propose a two-stage model for predicting enzyme function (EC number) from a protein sequence. First, we use semi-supervised learning to train a binary functional residue classifier with MSA pseudo-labels that quantifies evolutionary conservation from MSA data and the mCSA database as true functional residue labels. The binary classifier is used to filter non-functional residues and generate a fixed size representation from the predicted functional residue embeddings, which we call function-aware protein representations. Using function-aware protein representations, we train an EC number classification model. In this section, we provide an overview of the two-stage model and a detailed explanation of the training objectives for each stage.

#### Overview of the Two-stage Pipeline

As demonstrated in Figure 2, we propose a two-stage training pipeline for the EC number classification task. It is well established within the evolutionary biology community that a protein’s function is executed by a select few functional residues and the primary goal of the structure is to provide a stable scaffold that correctly positions these residues in 3D space [7, 50]. Thus, our approach is based on identifying these functional residues in order to remove the remaining residues and generate a function-aware representation. Previous studies [*e*.*g*., 10, 11, 49] attempt to identify functional residues via self-supervised learning where labels are generated from the MSAs. In stage one of our work, we propose to expand experimentally characterized functional datasets with MSA-priors via semi-supervised learning to train a binary classifier. To generate our function-aware representation, we utilize the confidence scores from the binary classifier to weight the hidden representation of the top-k most confident residues. In stage two of our work, we employ a simple MLP network to decode the function-aware representation into an EC number prediction, which we train via supervised learning.

**Figure 2.**
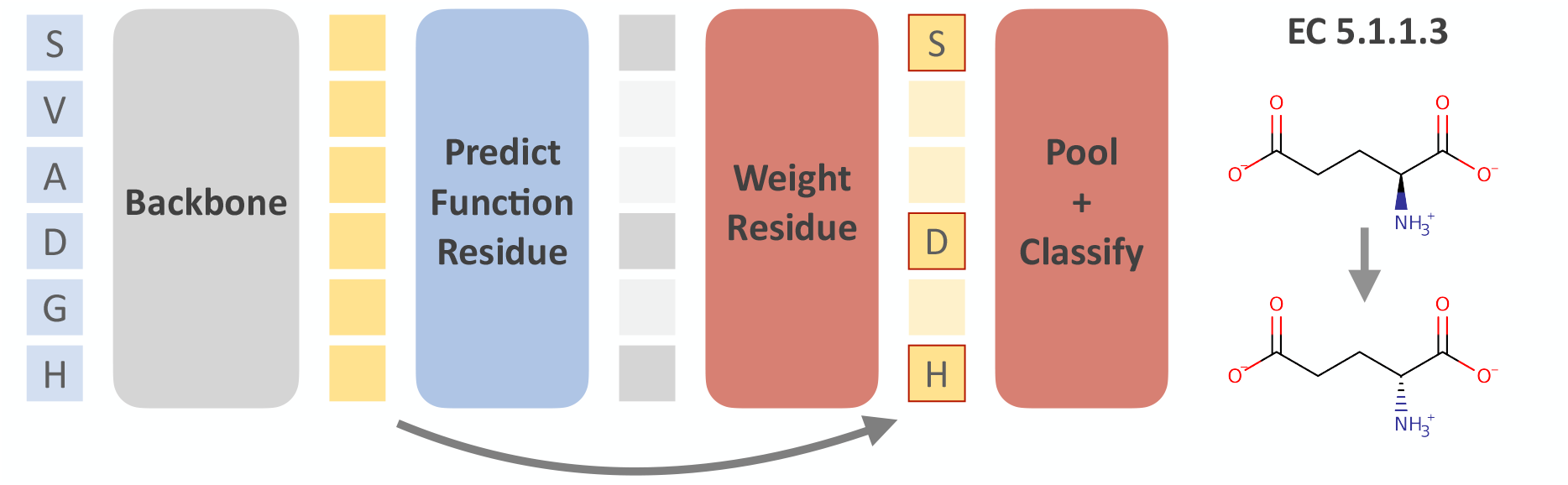
Overview of our two-stage EC number classification framework. In the first stage, our model predicts which residues are likely functional. In the second stage, it pools these predicted functional residues to obtain a fixed-length function-aware representation, which is decoded into an enzyme commission (EC) number prediction using an MLP.

#### Stage 1: Functional Residue Identification with MSAs

We start by describing the functional residue classification model training. Denote the protein sequence as 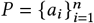 where *a*_*i*_ ∈ 𝒜 indicates the amino acid at index *i*, and 𝒜 is the set of 20 amino acids. In practice, we utilize the ESM model to extract representations, which can be substituted with those obtained from AlphaFold or other backbones. We note the embedding by {*e*_1_, ···, *e*_*n*_} = ESM *P*, where *e*_*i*_ denotes the contextualized hidden representation of *a*_*i*_ extracted from the pre-trained protein language model ESM [24, 36]. With the M-CSA (Mechanism and Catalytic Site Atlas) dataset, we have labeled data for every *a*_*i*_ indicating whether this amino acid is functional. We denote the labelled dataset 𝒟^sup^ = {*e*_*i*_, *y*_*i*_} where *y*_*i*_ ∈ {0, 1} is a binary label and we collect all amnio acid from all proteins.

In addition to using labeled data from the mCSA database, our functional residue dataset is further augmented with pseudo-labels generated from multiple sequence alignments (MSAs) of CATH v4.3 [42] (see Figure 3). To pseudo-label residues, we calculate the entropy of the following probability distribution derived from a protein’s MSA as a proxy of evolutionary conservation of a site *a*_*i*_ :

**Figure 3.**
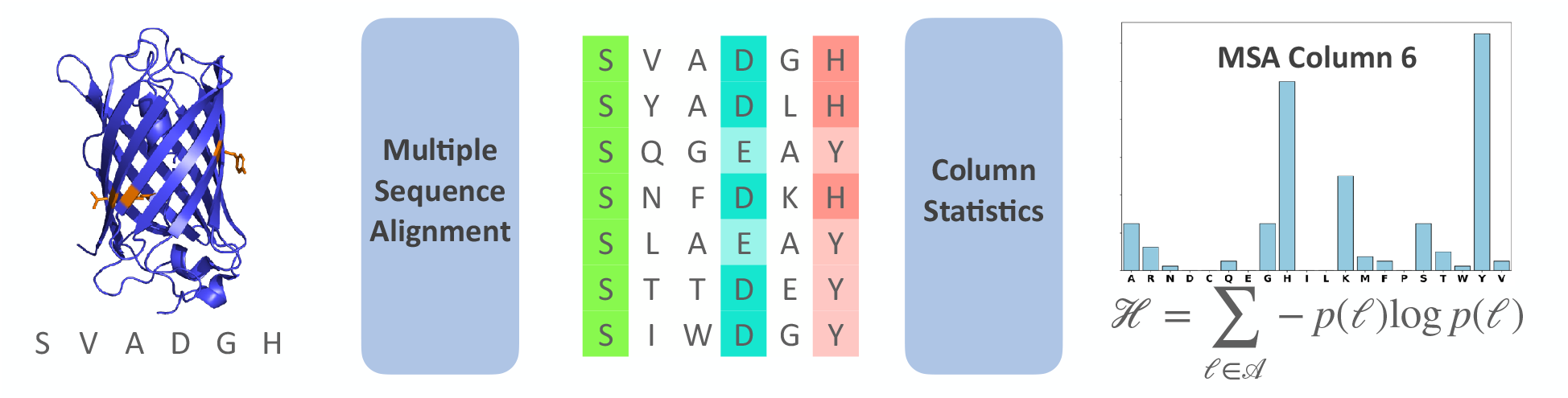
Functional residue pseudo-label generation from multiple sequence alignments. We introduce a self-supervised technique for labeling functionally relevant residues in a protein sequence from its MSA. More specifically, conserved positions— the 10% of columns with the lowest entropy—are labeled as functional.

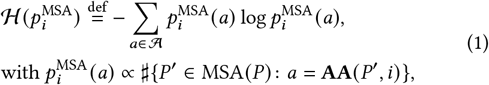

where *a* ∈ 𝒜 is one of the 20 amino acids, **AA** is the function that outputs the amino acid type at position *i*, MSA *P* denotes the set of sequences that are best aligned with *P* via multiple sequence alignment on UniRef90 [48], and ♯{·} denotes the number of elements of a set, and ℋ (*p*) is the entropy function of distribution *p*·

In practice, we calculate the entropy for all the positions *a*_*i*_ in one protein sequence, and pseudo-label the top 10% amino acids with the lowest entropy as functional residues, and we pseudo-label the MSA labeled dataset as 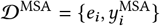, where pseudo-label 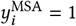 is positive for selected 10% sites and 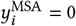 for the 90% remaining sets. Because the size of the negative class 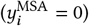 is 9 times of that of the positive class, we resample the negative class to make the two classes balanced.

To validate the reliability of the pseudo-labels, we use the MCSA labeled dataset as a case study. By extracting all MSA files, computing entropy scores, and designating the lowest 10% entropy residues in each sequence as functional, while considering the rest as non-functional, we achieve a recall of 0.5, precision 0.1 and F1 score 0.1, while random prediction baseline gets recall 0.4, precision 0.01, F1 0.02. This outcome suggests that this straightforward criterion of a column in the MSA having low-entropy effectively identifies functional residues within a protein sequence. Additional findings and outcomes are detailed in Section 4.

With 𝒟^sup^ and 𝒟^MSA^, one for labeled dataset and the other for the pseudo-label dataset, the loss function consists of two logistic loss ℒ terms. We propose to train the binary classifier *f* with the following confidence-aware training objective,

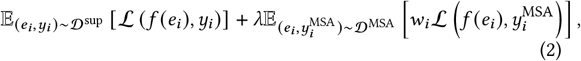

which is the sum of the standard cross entropy loss ℒ_*CE*_ on the MCSA dataset 𝒟^sup^, and that of the MSA pseudo-labeled 𝒟^MSA^ weighted by additional confidence threshold *w*_*i*_ :

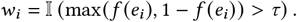

Here we assume that *f* (*e*_*i*_) ∈ [0, 1] is the output of the sigmoid gate, and *τ* is a scalar hyperparameter which controls the confidence threshold *w*_*i*_. In practice, we set *τ* = 0.9 and *λ* = 1.

The intuition of the threshold *w*_*i*_ is to ensure that, for each data instance, our trained model exhibits high confidence on the pseudo-label that is used for training. If the confidence level for a particular data instance is low, we discount the corresponding pseudo-label during gradient computation. Similar strategies have been employed in the past for disregarding low-confidence predictions in semi-supervised learning [44], domain adaptation [15], and other related problems where researchers utilize pseudo-labels. Doing this allows us to select the pseudo-labels that aligns with the supervised dataset during training, thereby enhancing the model’s robustness and generalization ability: the pseudo-labels that exhibit large disagreement with the MCSA labels should be discarded to avoid potential harmful bias.

#### Stage 2a: Function-Aware Protein Representation

Given a learned model *f*(·), we construct a representation of each protein *P* by aggregation the information from their functional residuals as predicted by *f* (·).

Let *P* = {*a*_*i*_, ···, *a*_*n*_} be a protein with sequence length *n*, and the ESM embedding *e*_*i*_ of each amino acid (recall these embeddings are contextualized by other amino acids in the sequence). We construct the function-aware representation of protein *P* by a weighted sum of *e*_*i*_ :

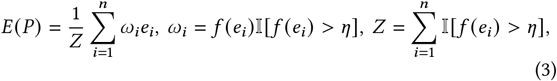

where each *e*_*i*_ is weighted by *ω*_*i*_ which is a thresholded variant of *f* (*e*_*i*_) that predicts whether *e*_*i*_ is functional. The threshold *η* is a hyperparameter dictating the number *Z* of amino acids considered in computing the protein embedding. Instead of employing uniformly weighted average pooling or learnable attention layers, this approach leverages labeled functional residue data to directly obtain the protein embedding based on its functional residues. In practice, we set *η* to 80% quantile for one given protein sequence so that *Z* = 80% × *n*.

#### Stage *2b: Functional Annotation Classification*

Using the function-aware protein representations in Eqn (3), we train a classification model *g* for the EC number classification task. We encode each 4-digit EC number into one class, and train a 4096-way classification head on the EC number dataset. We denote the dataset 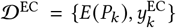, where *k* indexes the proteins in the dataset. Denote ℒ_CE_ as cross entropy loss, we use the weighted cross entropy loss as the training objective,

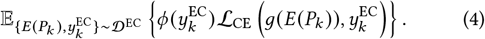

Here, *N* represents the number of EC classes, *ϕ*(·) weights each protein *P*_*k*_ based on its EC class 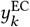. In practice, we set *ϕ* as inverse the frequency of each class in the training set. The weight *ϕ*(·) is needed due to the imbalanced nature of the EC number classes, and we observe that it establishes a baseline comparable to the state-of-the-art method CLEAN [54], while reducing the training cost by ∼80%. Compare to CLEAN’s simple MLP network, we choose *g* to be a three-layer MLP network, with layer normalization layers and a skip connection for the second layer.

## 4 Results

### 4.1 Experiment Details

#### Dataset

We summarize the datasets used in this paper in Table 2. We first introduce the functional residue identification datasets and MSA datasets we use, and then summarize the EC number classification training and test set details.

**Table 2:** Datasets to train and evaluate the two stages of our framework. For each protein in the mCSA dataset, we label the annotated residues as positive and all other residues as negative. Similarly, to generate the MSA training dataset, the top 10% of conserved residues of a protein are pseudo-labeled as positive, while the remaining 90% are labeled as negative.

| Dataset | Classification Task | Number of Proteins | Number of Classes |
| --- | --- | --- | --- |
| MSA | Functional Residue | 19,000 | 2 |
| mCSA train [34] | Functional Residue | 403 | 2 |
| mCSA val [34] | Functional Residue | 417 | 2 |
| CLEAN (100% split) [54] | EC Number | 220,000 | 4,096 |
| New-392 [54] | EC Number | 392 | 177 |
| Price-149 [32, 39] | EC Number | 149 | 55 |

For supervised learning of the functional residues in a protein sequence, we use the Mechanism and Catalytic Site Atlas (mCSA [34]), a database of 820 protein sequences with 3716 annotated functional residues. While mCSA provides mechanistic roles for the functional residues (e.g. general acid, transition-state stabilization, etc), we curate our dataset with functional residues as positive class and the remaining residues as the negative class. To evaluate our ability to identify functional residues, we split mCSA into a training set of 403 protein sequences with 1818 annotated residues and a validation set of 417 protein sequences with 1898 annotated residues. To limit homology-based data leakage, validation set proteins are at most 50% sequence similar to those in the training set.

For the EC number classification task, we use the training and test sets from CLEAN [54]. Briefly, the training set consist of 220*K* enzyme sequences by clustering and subsampling SwissProt [4], the human-annotated subset of UniProtKB. For evaluation, we use the New-392 and `Price-149` test sets. The `Price-149` dataset, introduced by ProteInfer [39], consists of 149 proteins [32] across 55 distinct EC numbers, including eight out-of-distribution (OOD) EC classes with respect to our training set. The New-392 dataset, introduced by CLEAN [54], is a more comprehensive benchmark, featuring 392 enzyme sequences distributed across 177 distinct EC numbers curated from Swiss-Prot. Additionally, we evaluate the impact of removing homologous proteins from the training set with respect to the `Price-149` and `New-392` test sets at various sequence identity cutoffs (*e*.*g*., 10%, 30%, 50%, 70%). These filtered training sets were originally introduced in CLEAN and obtained from their repository.

#### Hyperparameter Configurations

Our EC framework uses two-stage training procedure. In the first stage, we pre-generate ESM2 embeddings and train a residue-level binary classifier to identify functional residues using semi-supervised learning, where we use the mCSA dataset as positive true labels and low-entropy residues in the MSA dataset as positive pseudo-labels. The model is a two-layer MLP with 256 hidden dimension, and is trained with 10^™5^ learning rate, 0.01 weight decay, 8192 tokens in one iteration, and 10, 000 iterations with the AdamW optimizer. In the second stage, we train a EC number classifier. This model is a three-layer MLP network with LayerNorm layers and a skip connection for the second layer, 4096 hidden dimension, 4096 batch size, 10^™5^ learning rate, 0.0 weight decay, and is trained for 10, 000 iterations using the AdamW optimizer. During training, we make use of label smoothing (0.5) and mix-up data augmentation (*α* = 0.2). Using one A-40 GPU, pre-generating the the ESM2 embedding requires ∼30 minutes, training the functional residue binary classifier requires ∼15 minutes, and training the EC classifier requires ∼10 minutes.

#### Baselines

We compare our model against a suite of computational and machine learning-based baselines. Homology search has been the de-facto method to infer the function of a target protein lacking experimental validation. BLASTp [1] retrieves previous labeled enzymes in the training set with high sequence similarity. These homology search methods then assigns the retrieved function as the predicted function for the uncharacterized protein. We implement this baseline following ProteInfer [39]. Moreover, we compare to several machine learning-based baseline algorithms. DeepEC [38], EnzBERT [5] and ECPred [9] are classification frameworks for predicting the EC number. CLEAN [55] is trained with contrastive learning where proteins with the same EC number are considered as positive pairs and pre-generated EC number cluster representation are used to conduct classifcation during inference.

### 4.2 Enzyme Commission (EC) Number Classification

Here, we evaluate the second training stage: EC number classification using a function-aware protein representations. We first demonstrate the results with 100% sequence similarity data splits with respect to the `New-392` and `Price-149` datasets (Table 3). In the first block, We report the traditional sequence alignment baseline, BLAST. In the middle block, we report literature machine learning models, such as ECPred [9], EnzBERT [5], DeepEC [38], ProteinInfer [39], and the recent SOTA method CLEAN [54]. The last block demonstrates our results.

**Table 3:**
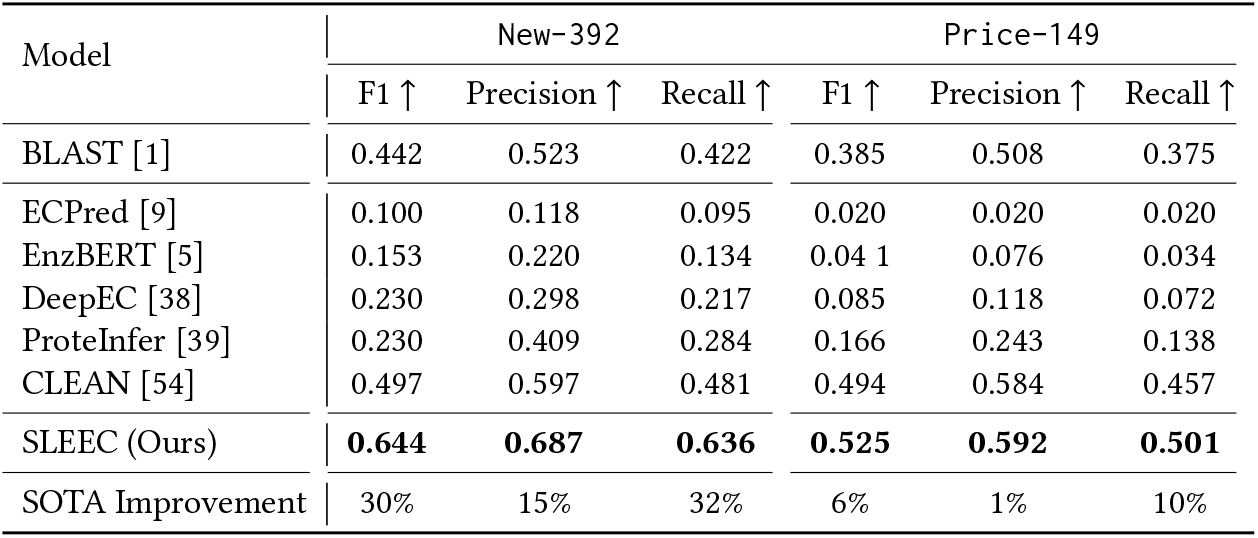
Classification performance comparisons on the New-392 and Price-149 EC number test sets. Top: Sequence homology-based approach. Middle: Literature machine learning frameworks. Bottom: Our framework. We report SLEEC classification metrics using the complete CLEAN training set (100% sequence similarity splits).

With respect to F1 score, SLEEC further improves over previous SOTA by 30% on `New-392` and 6% on `Price-149`. We note that improvements on price is much more challenging due to to 15% of the EC numbers in this test set lack training data and are out of distribution (OOD). To generalize to OOD EC number classes, we train an separate classification head to predict the three-digit EC number (247-class classifier). In order to evaluate without inadvertently leaking whether a sequence belongs to an OOD EC number, we create a new classification head. This head selectively utilizes the Softmax neurons from either the four-digit or the three-digit EC number classification heads, depending on the availability of training data for a specific four-digit EC class. This post-training classification head enables a comprehensive assessment of the model’s performance across various training dataset configurations, where the number of EC number classes lacking training data may vary. Additionally, it facilitates examination of the model’s robustness and generalization capabilities.

In Table 4, we compare the generalization capability of CLEAN and SLEEC using several homology-based data splits: 10%,30%, 50%, 70%. SLEEC outperforms CLEAN across the five data splits with an average F1 performance improvement of ∼35%. Notably, training SLEEC with the 50% sequence similarity filtered training set outperforms CLEAN trained with no homology filtering (100% dataset) by 18% (F1 0.584 vs 0.497).

**Table 4:** F1 scores on different homology-based data split settings. We evaluate performance on the New-392 and Price-149 test sets when trained at different homology-filtering thresholds. These homology-filtered datasets are provided by CLEAN.

| Test Dataset | New-392 |  |  |  |  | Price-149 |  |  |  |  |
| --- | --- | --- | --- | --- | --- | --- | --- | --- | --- | --- |
| Dataset Size (K) | 10% | 30% | 50% | 70% | 100% | 10% | 30% | 50% | 70% | 100% |
|  | 7.8 | 9.8 | 29.4 | 74.5 | 22.7 | 7.8 | 9.8 | 29.4 | 74.5 | 22.7 |
| CLEAN | 0.312 | 0.362 | 0.394 | 0.416 | 0.497 | 0.235 | 0.267 | 0.304 | 0.372 | 0.495 |
| SLEEC (Ours) | **0.402** | **0.447** | **0.584** | **0.605** | **0.644** | **0.247** | **0.284** | **0.328** | **0.399** | **0.525** |
| Improvement | 29% | 23% | 48% | 45% | 30% | 5% | 6% | 8% | 7% | 6% |

**Table 5:** Robustness comparisons on the New-392 and Price-149 test sets using bio-based sequence modifications. We report F1 scores under various sequence modification settings. ‘Sensitivity’ demonstrates the difference between the results on the original test set and on the augmented test set as a percentage. ‘Improvement’ demonstrates the difference in F1 performance between CLEAN and SLEEC. We use the same settings presented in Table 1.

| Model | New-392 |  |  |  | Price-149 |  |  |  |
| --- | --- | --- | --- | --- | --- | --- | --- | --- |
|  | Original | Halo-Tag | TAP-Tag | Ortholog | Original | Halo-Tag | TAP-Tag | Ortholog |
| CLEAN ↑ | 0.497 | 0.302 | 0.434 | 0.405 | 0.495 | 0.190 | 0.342 | 0.412 |
| CLEAN sensitivity ↓ | 0% | 39% | 13% | 19% | 0% | 62% | 31% | 17% |
| SLEEC (Ours) ↑ | **0.644** | **0.619** | **0.632** | **0.632** | **0.525** | **0.503** | **0.512** | **0.527** |
| SLEEC sensitivity ↓ | 0% | 4% | 2% | 2% | 0% | 4% | 3% | 0% |
| Improvement ↑ | 30% | 105% | 46% | 56% | 6% | 165% | 50% | 28% |

#### Robustness to Bio-based Sequence Modifications

In protein engineering workflows, protein sequences are frequently modified with functional tags or screened alongside homologs. Therefore, it is crucial for EC number predictions to be robust to such modifications, since they generally don’t alter enzymatic function. Building upon the case study presented in the introduction (Table 1), we compare the model performance of SLEEC vs CLEAN in these scenarios, using two N-terminus tags and orthologs with an ∼80% sequence similarity, as shown in Table 7. For all sequence modification experiments on `Price-149` and `New-392`, our method consistently exhibits superior robustness compare to CLEAN. We attribute this enhanced robustness to our function-aware representations, in contrast to CLEAN’s, which derives its representation through average pooling.

**Table 6:**
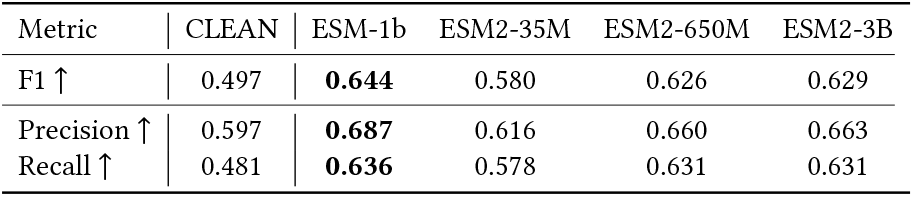
Metrics on the New-392 dataset using different back-bones. We evaluate the impact different ESM-based back-bones have on the EC number classification task.

| Metric | CLEAN | ESM-1b | ESM2-35M | ESM2-650M | ESM2-3B |
| --- | --- | --- | --- | --- | --- |
| F1 ↑ | 0.497 | **0.644** | 0.580 | 0.626 | 0.629 |
| Precision ↑ | 0.597 | **0.687** | 0.616 | 0.660 | 0.663 |
| Recall ↑ | 0.481 | **0.636** | 0.578 | 0.631 | 0.631 |

**Table 7:**
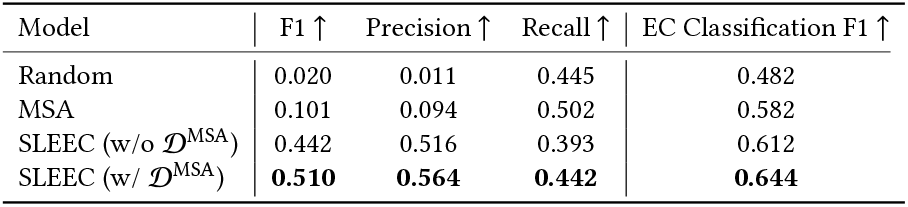
Functional residue prediction on mCSA validation set. We evaluate the binary classifier’s performance on identifying functional residues. MSA denotes applying low-entropy policy to distinguish functional residues. Here, we split the training and test set due to 50% sequence similarity to avoid data leakage. The last column provides downstream evaluations on the New-392 test set.

| Model | F1 ↑ | Precision ↑ | Recall ↑ | EC Classification F1 ↑ |
| --- | --- | --- | --- | --- |
| Random | 0.020 | 0.011 | 0.445 | 0.482 |
| MSA | 0.101 | 0.094 | 0.502 | 0.582 |
| SLEEC (w/o $\mathcal{D}^{\text{MSA}}$) | 0.442 | 0.516 | 0.393 | 0.612 |
| SLEEC (w/ $\mathcal{D}^{\text{MSA}}$) | **0.510** | **0.564** | **0.442** | **0.644** |

#### Backbone Ablations

In Table 6, we conduct an ablation study to compare the ESM-1b [36] backbone against three ESM2 backbones (*e*.*g*., 35M, 650M, 3B). ESM-1b serves as the backbone for both our literature transformer-based baseline studies and the backbone of SLEEC. Through ablation studies, we find that the specific choice of backbone has minimal impact, while the size of the backbone plays a more crucial role. ESM-1b is the backbone used by our literature transformer-based baseline studies and is the backbone SLEEC is built with. From these ablations studies, we observe that the backbone does not have a significant impact but rather the size of the backbone is much more important. Once we apply a larger backbone, *e*.*g*., ESM2 3B, ESM2 650M, the fluctuation in performance is not significant and can likely be mitigated through hyperparameter tuning. Remarkably, despite the ESM2 35M backbone being 18.6 times smaller than CLEAN’s backbone, SLEEC still achieves an F1 score that surpasses CLEAN by ∼17% (0.58 vs 0.497). This highlights the effectiveness of learning function-aware protein representations.

#### Impact of Classification Head vs. Pooling

To disentangle the performance gains attributed to the multi-label classification head versus the functional residue pooling, we performed a componentwise analysis. Results on the New-392 dataset show that replacing CLEAN’s contrastive loss with a multi-label classification head accounts for part of the improvement, raising the F1 score from 0.49 to 0.55. However, the integration of our functional residue pooling strategy provides a more substantial boost, further increasing the F1 score to **0.64**. On the Price-149 dataset, the classification head alone yields results comparable to the baseline (0.49), whereas the full SLEEC model (with pooling) achieves 0.53. This confirms that while the classification head improves training efficiency, the core performance gain and generalization capability primarily stem from our proposed functional residue pooling module.

#### Sensitivity to Functional Residue Ratio

To validate our assumption that enzyme function is determined by a sparse set of residues, we conducted an ablation study on the selection threshold *η* (the percentage of residues pooled for representation) on the New-392 dataset. As shown in Table 8, the model maintains robust performance (F1 ≈ 0.64) when retaining only 5% to 20% of the residues. This confirms that SLEEC effectively captures the sparse, critical functional signals. Notably, increasing the ratio to 50% introduces irrelevant background noise, leading to a slight performance drop (0.62).

**Table 8:** Ablation study on the percentage of functional residues used for pooling (tested on New-392).

| Residue Ratio ($\eta$) | 5% | 10% | 20% | 50% |
| --- | --- | --- | --- | --- |
| **SLEEC (F1)** | 0.64 | 0.64 | 0.64 | 0.62 |

### 4.3 Evaluation on The ProteInfer Test Set

To further demonstrate the robustness and generalization capability of SLEEC, we evaluated our model on the **ProteInfer** benchmark datasets [39]. This benchmark includes both Enzyme Commission (EC) number prediction and Gene Ontology (GO) term classification tasks. We adopted the same hyperparameter settings as CLEAN for a fair comparison. As shown in Table 9, SLEEC consistently outperforms CLEAN on both tasks. Specifically, on the EC split, SLEEC achieves an F1 score of **0.95** (vs 0.92), and on the GO split, it reaches **0.86** (vs 0.80). These results confirm that our function-aware representation learning framework generalizes effectively across different functional annotation ontologies and data splits.

**Table 9:** Evaluation on The ProteInfer Test Set.

| Method | EC (F1) | GO (F1) |
| --- | --- | --- |
| CLEAN | 0.92 | 0.80 |
| **SLEEC (Ours)** | **0.95** | **0.86** |

### 4.4 Functional Residue Classification

In this section, we assess the initial training stage, which involves a binary classifier aimed at identifying functional residues within a protein sequence via semi-supervised learning. For this evaluation, we split the mCSA dataset into a training and validation set using a 50% sequence similarity threshold. We report binary classification metrics of F1, Precision and Recall in Table 7. We compare against the following baselines: a random classifier, our low-entropy MSA policy for pseudo-labeling residues, and supervised learning with the mCSA training set. The low-entropy MSA baseline shows that MSA conservation alone is inadequate for accurately distinguishing functional residues from structurally conserved positions, resulting in low precision. However, when combined with mCSA for semi-supervised learning, it boosts the F1 score from 0.44 to 0.51, marking a 15.4% improvement, and enhances the downstream EC classification F1 score by 5.2%. This underscores the effectiveness of our representations in improving performance on downstream functional tasks.

## 5 Limitations and Ethical Considerations

In this work, we set up our experiments based on the previous SOTA model, CLEAN, in which the backbone model weights are frozen. In the future, we will test more training settings, such as sparse fine-tuning and memory efficient optimizers. We currently compare our method with existing benchmarks and plan to include additional wet-lab comparisons in future work. Our method is currently limited to substrates with defined EC numbers. However, generalization of our framework to new substrates has the potential to significantly enhance the identification of enzymes that enable sustainable procurement of agricultural and chemical commodities, pharmaceuticals, and food ingredients.

## 6 Conclusions

In this work, we present a residue-level semi-supervised learning technique to learn function-aware protein representations and demonstrate their capability to improve the downstream EC number classification task. Additionally, our function-aware representations improve robustness to routine sequence modifications encountered in protein engineering workflows and provide residue-level interpretable predictions. Moving forward, we plan to further refine our MSA-based pseudo-labeling algorithm to improve the precision of pseudo-labeled functional residues and extend the classification framework to reactions lacking EC numbers and enzymes with promiscuous catalytic activity. Finally, we believe our proposed function-aware protein representations can be generalized to additional downstream functional tasks that are also evolutionarily encoded.

